# Superior Efficacy of Combination Antibiotic Therapy versus Monotherapy in a Mouse Model of Lyme Disease

**DOI:** 10.1101/2023.04.25.538305

**Authors:** Yasir Alruwaili, Mary B. Jacobs, Nicole R. Hasenkampf, Amanda C. Tardo, Celine E. McDaniel, Monica E. Embers

**Affiliations:** Department of Clinical Laboratory Sciences, College of Applied Medical Sciences, Jouf University, Sakaka, 72388, Saudi Arabia; Division of Immunology, Tulane National Primate Research Center, Tulane University, Covington, Louisiana 70433, USA; Department of Tropical Medicine, School of Public Health and Tropical Medicine, Tulane University, New Orleans, Louisiana 70112, USA

## Abstract

Lyme disease (LD) results from the most prevalent tick-borne infection in North America, with over 476,000 estimated cases annually. The disease is caused by *Borrelia burgdorferi (Bb)* which transmits through the bite of Ixodid ticks. Most cases treated soon after infection are resolved by a short course of oral antibiotics. However, 10-20% of patients experience chronic symptoms because of delayed or incomplete treatment, a condition called Post-Treatment Lyme Disease (PTLD). Some *Bb* persists in PTLD patients after the initial course of antibiotics and an effective treatment to eradicate the persistent *Bb* is needed. Other organisms that cause persistent infections, such as *M. tuberculosis*, are cleared using a combination of therapies rather than monotherapy. A group of Food and Drug Administration (FDA)-approved drugs efficacious against *Bb* were used in monotherapy or in combination in mice infected with *Bb*. Different methods of detection were used to assess the efficacy of the treatments in the infected mice including culture, xenodiagnosis, and molecular techniques. None of the monotherapies eradicated persistent *Bb*. However, 4 dual combinations (doxycycline + ceftriaxone, dapsone + rifampicin, dapsone + clofazimine, doxycycline + cefotaxime) and 3 triple combinations (doxycycline + ceftriaxone+ carbomycin, doxycycline + cefotaxime+ loratadine, dapsone+ rifampicin+ clofazimine) eradicated persistent *Bb* infections. These results suggest that combination therapy should be investigated in preclinical studies for treating human Lyme disease.

## Introduction

Lyme disease (LD), caused by the *Borrelia burgdorferi* (*Bb*) sensu lato complex, is the most common tick-borne illness in the United States with more than 476,000 cases annually (1). Approximately 10-20% of LD patients further develop post-treatment Lyme disease (PTLD) and continue to experience symptoms for 6 months or more following treatment. Persistent *Bb* infection that is not eradicated by guideline-adherent antibiotic treatment is strongly suspected to contribute to PTLD, although there are other proposed causes as well (2–5).

The Infectious Disease Society of America (IDSA) and other international agencies of infectious diseases recommend monotherapy (single drug treatment) with doxycycline or amoxicillin to treat LD (1, 2). Monotherapy can be effective 3 to 30 days after the tick bite with response rates of 89% or higher (3, 4). However, treatment delays significantly affect antibiotic efficacy, and many patients experience chronic musculoskeletal pain, fatigue, and cognitive dysfunction (5-8). Studies on patients with PTLD show not only that immune activation/inflammation lingers in the brain, but also that dynamic changes in the frontal lobe associated with slower processing speeds occur (9, 10). Treatment for PTLD is mired by a lack of tests for persistent infection and mixed results on treating chronic Lyme disease in clinical trials (11-14).

Studies in mice, dogs, and nonhuman primates have demonstrated persistence of the spirochetes after doxycycline or ceftriaxone treatment (15-21). In vitro studies have demonstrated improved efficacy with combinations of antibiotics (22-24). The currently available studies that address these limitations demonstrate that a combination of daptomycin + cefoperazone + doxycycline eradicated persistent Bb in culture and a combination of daptomycin+ doxycycline+ ceftriaxone eradicated persistent Bb in mice (25, 26).

Combination therapy is an alternative approach to treat *Bb* infection and is already well-established for treatment of persistent bacteria like *M. tuberculosis* and *Brucella melitensis* in humans and animal models (27-31). Only one randomized clinical trial that used more than one drug (doxycycline and ceftriaxone) has been performed, but the drugs were administered at separate time points rather than concurrently and treatment failed to demonstrate patient improvement (11). A separate non-randomized study used a combination of clarithromycin and hydroxychloroquine. This combination was as effective as prolonged tetracycline treatment but the inclusion criteria were not clearly defined, and the study was non-randomized (12). Barriers to combination therapy clinical trials include disagreement on the cause of PTLD and limited preclinical evidence for efficacy.

The study described here used a panel of drugs as monotherapy and, separately, a panel of combination therapies to treat long-term (persistent) *Bb* infection. The presence of persistent *Bb* infection following treatment was determined using multiple methods of detection. These results provide data to further demonstrate *Bb* persistence following monotherapy and the potential for combination therapy to prevent persistent *Bb* infection. These results also demonstrate feasibility of combination therapy as an alternative treatment regimen to monotherapy for disseminated infection that can lead to PTLD.

## Results

### Confirmation of infection in the monotherapy treatment arm

The experimental procedure for testing efficacy of drugs and combinations of drugs is depicted in **Figure 1**. Infection with *Bb* was confirmed by either an ear punch culture positive for *Bb* or a 5-antigen serology result positive for 2 or more *Bb* antigens at days 21 or 60. Mice were considered infected if either test was positive. Twenty out of 50 (40%) mice had positive ear punch results while 50 out of 50 (100%) had positive 5-antigen test results. Table S1 shows the results of each group.

**Figure 1.**
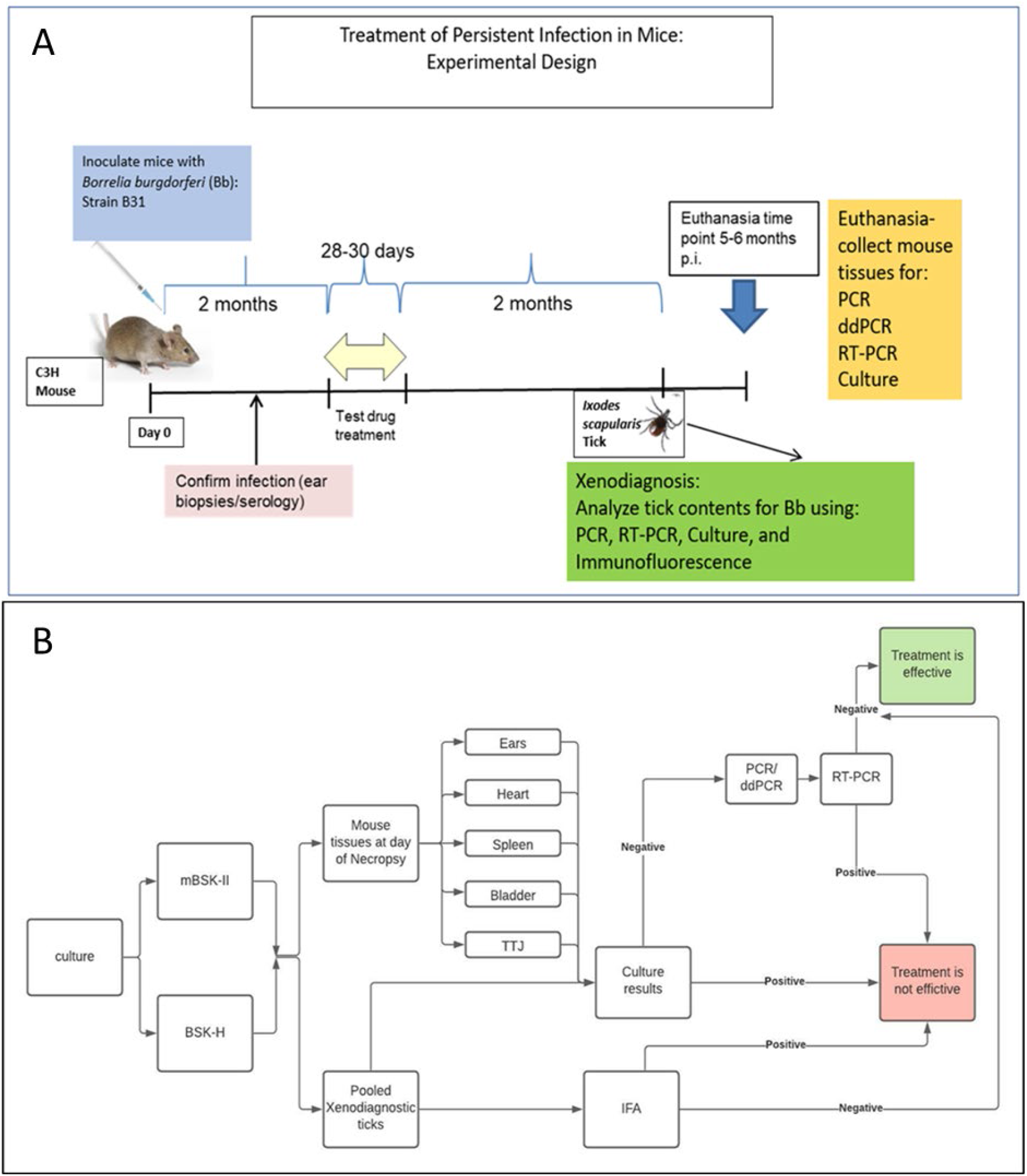
Experimental protocol for assessment of efficacy in mice. (A) Mice were infected and kept for 2 months prior to treatment. After the treatment, 2 more months elapsed before determination of persistent infection, including xenodiagnoses, culture, and molecular detection. (B) Flow chart for the determination of efficacy.

### Determination of drug concentration in serum following treatment

Drugs that were be tested in mice are all FDA-approved and have been shown to kill or prevent the growth of *Borrelia burgdorferi* in different growth phases or morphologies (i.e. sessile, round-body forms, and aggregates). These compounds have shown promise *in vitro* but had not been tested in an animal model of persistent infection (24, 25, 32-36). Mice with confirmed infections were treated with a single drug. Each drug was administered at the highest non-toxic concentration. (**Table 1**) The drug concentration in the serum was calculated using the modified Kirby Bauer assay (MKB). One drug per route of administration (rifampicin, OG; cefotaxime, IP; carbomycin SC) was selected to measure serum concentrations at 24-hour post-treatment during weeks 1-3 of treatment.

**Table 1:** Concentrations, doses, routes, and number of mice.

| Treatment | Concentration | Dose<br>(1x per day) | Route<br>(OG, IP, SC) | n |
| --- | --- | --- | --- | --- |
| **Controls** |  |  |  |  |
| Infected, untreated<br>(OG Ctrl) | Water | 0.2 ml/mouse | OG | 3 |
| Infected, untreated<br>(IP Ctrl) | PBS | 0.2 ml/mouse | IP | 5 |
| Infected, untreated<br>(SC Ctrl) | 0.09% Saline +5%<br>DMSO | 0.2 ml/mouse | SC | 2 |
| **Total** |  |  |  | 10 |
| **Therapies** |  |  |  |  |
| Azlocillin | 15 mg/mL | 100 mg/kg | IP | 5 |
| Bactrim | 10 mg/mL | 62.5 mg/kg | OG | 5 |
| Disulfiram | 20 mg/mL | 100 mg/kg | OG | 5 |
| Carbomycin | 25 mg/mL | 200 mg/kg | SC | 5 |
| Dapsone | 5 mg/mL | 15 mg/kg | OG | 5 |
| Rifampicin | 10 mg/mL | 30 mg/kg | OG | 5 |
| Cefotaxime | 25 mg/mL | 100 µg/mouse | IP | 5 |
| Loratadine | 5 mg/mL | 20 mg/kg | OG | 5 |
| **Total** |  |  |  | 40 |

OG, oral gavage; IP, intraperitoneal; SC, subcutaneous

The average concentration of rifampicin 24-hours post-treatment for weeks 1-3 was 2.588 µg/mL (range= 1.23 – 3.87) with a total of 2 pooled serum samples for every week. At day 66 (week 1), the average concentration of cefotaxime at the trough level was calculated as 2.454 ug/ mL (range 0 - 5.18) of 2 pooled sera. At day 73 (2nd week of treatment), the average concentration of carbomycin at the trough level was 4.8 ug/ mL (range 4.42 - 5.2 ug/ mL) of 2 pooled sera. Supplementary materials (Tables S2-5 and Figures S1-3) include more details of MKB results.

These acquired concentrations were higher than the MIC of these drugs in previous studies. The MIC of rifampicin against persisting *Bb* is ≤0.4 μg/mL (33). The MIC and MBC of cefotaxime against persisting *Bb* are ≤0.03 μg/mL and ≤0.25 μg/mL, respectively (37). The MIC of carbomycin against persisting *Bb* is ≤0.25 μg/mL (32).

In addition, azlocillin and dapsone were selected to be tested via MKB but the interaction of these drugs, when tested with *S. aureus*, was limited to the concentration of 10 μg/ml and up for azlocillin, and 4100 μg/ml or more for dapsone. These minimum concentrations are not anticipated to be achieved in our mouse sera, precluding the use of this test for measuring antibiotic levels in blood.

### Culture of tissues from mice treated with monotherapy confirm Bb infection

Mice were divided into groups of 2-5 and administered either a test drug or control treatment. None of the monotherapy regimens completely eradicated *Bb*. The cefotaxime group had significantly fewer infected mice compared to the control (*P* = .0476), but *Bb* persisted in 1 mouse. The other monotherapy groups had 3-5 mice positive for *Bb* per group following treatment. There was no difference in the number of positive samples between BSK-H and mBSK-II media (**Table 2)**.

**Table 2:** Culture results of tissues in monotherapy.

| Treatments | Cultures of Tissues |  |  | P value |
| --- | --- | --- | --- | --- |
|  | BSK-H<sup>^</sup> | mBSK-II<sup>^</sup> | Combined Cultures<sup>^</sup> |  |
| Infected Untreated (OG) | 3/3 | ND | 3/3 |  |
| Infected Untreated (IP) | 5/5 | ND | 5/5 |  |
| Infected Untreated (SC) | 2/2 | ND | 2/2 |  |
| Azlocillin (IP) | 4/5 | ND | 4/5 | 1<sup>b</sup> |
| Bactrim (OG) | 5/5 | ND | 5/5 | 1<sup>a</sup> |
| Disulfiram (OG) | 5/5 | ND | 5/5 | 1<sup>a</sup> |
| Carbomycin (SC) | 5/5 | 5/5 | 5/5 | 1<sup>c</sup> |
| Dapsone (OG) | 3/5 | 3/5 | 3/5 | 0.4643<sup>a</sup> |
| Rifampicin (OG) | 3/5 | 3/5 | 3/5 | 0.4643<sup>a</sup> |
| Cefotaxime (IP) | 1/5 | 1/5 | 1/5 | 0.0467<sup>b*</sup> |
| Loratadine (OG) | 3/5 | 3/5 | 3/5 | 0.4643<sup>a</sup> |

ND, not done; OG, oral gavage; IP, intraperitoneal; SC, subcutaneous
Combined cultures compared to a= OG control group, b= IP control group, c= SC control group.
<sup>^</sup> Number of mice positive for at least one tissue/total number of mice tested.
\* Statistically significant. Fisher exact test, the result is significant at $p < .05$ .

### PCR and RT-PCR confirm *Bb* infection in and viability in tissues following monotherapy

*Bb* was detected by PCR using the *16S* primer to detect *Borrelia* species specific DNA in 2-5 mice from each monotherapy group. PCR using primers specific to the *ospC* gene in the B31 strain of *Bb* confirmed the presence of *Bb* in 6 out of 8 monotherapy groups. RT-PCR using the *16S* primers detected *Bb* RNA in at least 1 mouse in all groups, confirming the presence of viable *Bb* following monotherapy (**Table 3**).

**Table 3:**
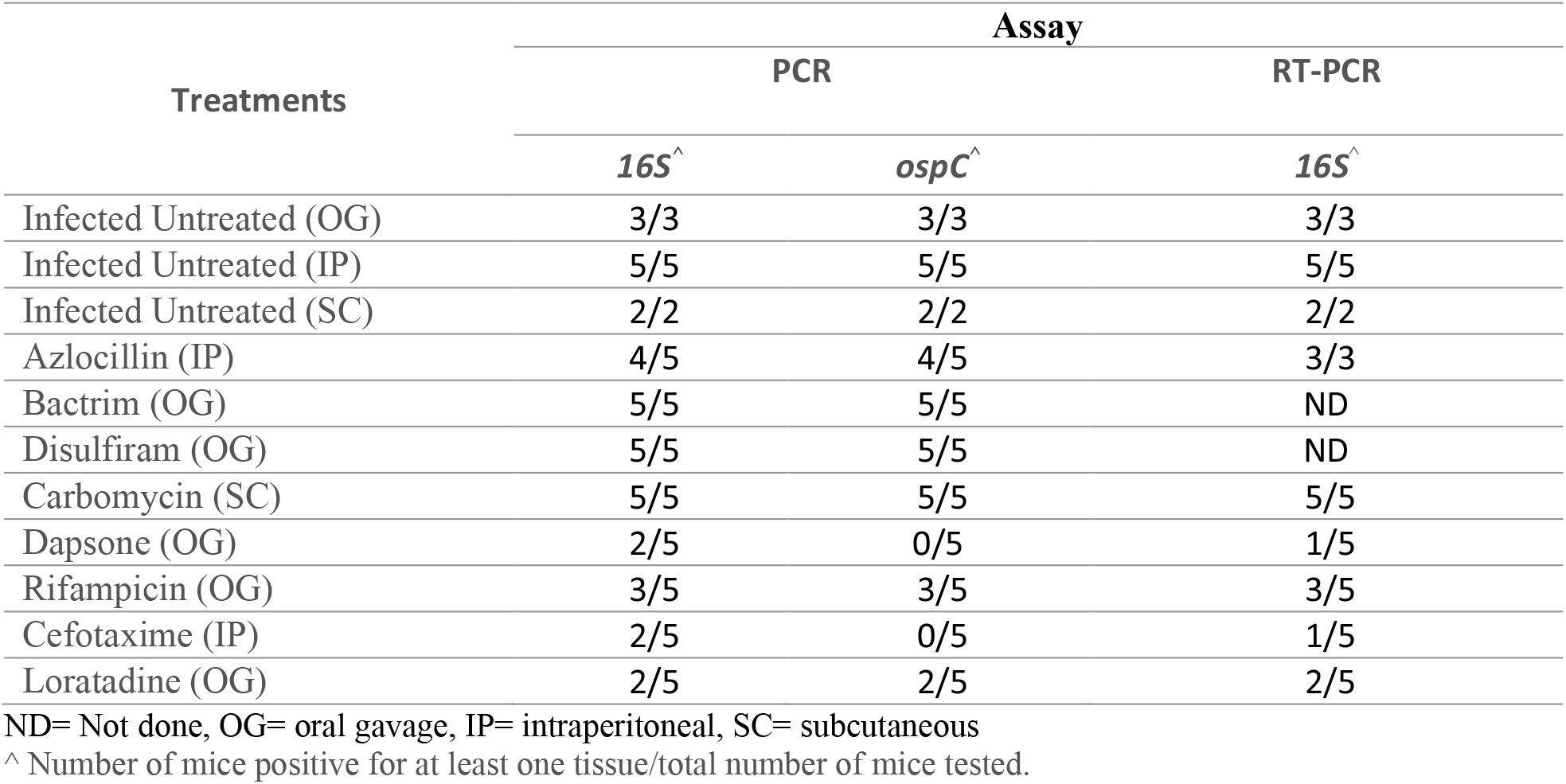
Molecular detection of Bb in Tissues of Mice.

### Cultured xenodiagnostic tick midguts harbor *Bb* following monotherapy

Cultured tick midguts identified Bb infection in at least 1 mouse from each treatment group. The cefotaxime group had significantly fewer infected mice compared to the control (*P* = .0476), but *Bb* persisted in 1 mouse. The other monotherapy groups had 2-5 mice positive for *Bb* per group following treatment. The control mice had a 70% detected rate of infection.

BSK-H or mBSK-II were used to culture the tick midguts. The results were identical for BSK-H and mBSK-II culture media with the exception of the group treated with dapsone for which *Bb* was detected in 40% of tick midguts cultured in BSK-H media and in 60% of tick midguts cultured in mBSK-II (**Table 4**). Results from both media conditions were combined for the analysis.

**Table 4:** Culture results for xenodiagnostic ticks following monotherapy treatment of infected mice.

| Treatments | Cultures of Xenodiagnostic Ticks |  |  | P value |
| --- | --- | --- | --- | --- |
|  | BSK-H<sup>^</sup> | mBSK-II<sup>^</sup> | Combined Cultures<sup>^</sup> |  |
| Infected Untreated (OG) | 1/3 | ND | 1/3 |  |
| Infected Untreated (IP) | 5/5 | 3/3 | 5/5 |  |
| Infected Untreated (SC) | 1/2 | 1/2 | 1/2 |  |
| Azlocillin (IP) | 4/5 | ND | 4/5 | 1<sup>b</sup> |
| Bactrim (OG) | 5/5 | ND | 5/5 | 0.1071<sup>a</sup> |
| Disulfiram (OG) | 3/5 | ND | 3/5 | 1<sup>a</sup> |
| Carbomycin (SC) | 4/5 | 4/5 | 4/5 | 0.5238<sup>c</sup> |
| Dapsone (OG) | 2/5 | 3/5 | 3/5 | 1<sup>a</sup> |
| Rifampicin (OG) | 2/5 | 2/5 | 2/5 | 1<sup>a</sup> |
| Cefotaxime (IP) | 1/5 | 1/5 | 1/5 | 0.0467<sup>b*</sup> |
| Loratadine (OG) | 3/5 | 3/5 | 3/5 | 1<sup>a</sup> |

ND= Not done, OG= oral gavage, IP= intraperitoneal, SC= subcutaneous
Combined cultures compared to a= OG control group, b= IP control group, c= SC control group
<sup>^</sup> Number of pooled ticks per mouse positive / total number of mice tested.
\* Statistically significant. Fisher exact test, the result is significant at $p < .05$ .

### PCR and RT-PCR confirm *Bb* presence in and viability in xenodiagnostic ticks following monotherapy

*Bb* was detected by PCR using the *16S* and *ospA* primers in 1-5 tick midgut samples in all but 1 monotherapy group (**Table S5**). The primer for *ospA* did not detect *Bb* in the rifampicin group. Control groups from both PCR assays detected *Bb* in 55% of samples. In particular, *Bb* was not detected by PCR in the group that received the placebo by subcutaneous injection. Bb was detected by RT-PCR using the *16S* and *ospA* primers in 1-5 tick midgut samples in all monotherapy groups. Control groups from both PCR assays detected *Bb* in 95% of samples.

### Confirmation of infection in combination therapy

Infection with *Bb* was confirmed in the same way as animals that received monotherapy: by either an ear punch culture positive for *Bb* or a 5-antigen serology result positive for 2 or more *Bb* antigens at days 21 or 60. Mice were considered infected if either test was positive. Twenty-seven out of 65 (42%) mice had positive ear punch results while 65 out of 65 (100%) had positive 5-antigen test results. **Table S6** shows the results of each group.

### Culture of tissues from mice treated with combination therapy confirm *Bb* eradication

Combination therapy was efficient at clearing *Bb* infection depending on the series of drugs used. Mice were divided into groups of 2-5 and administered either a combination of test drugs or control treatment. Eight out of 11 groups receiving a combination of drugs were 0% positive for *Bb* infection, a significant reduction compared to the control groups (*P* =.05). The groups positive for Bb infection were azlocillin+ Bactrim (100%), disulfiram+ Bactrim (100%), and disulfiram + azlocillin (80%). There was no difference in the number of positive samples between BSK-H and mBSK-II media (**Table 5**).

**Table 5:**
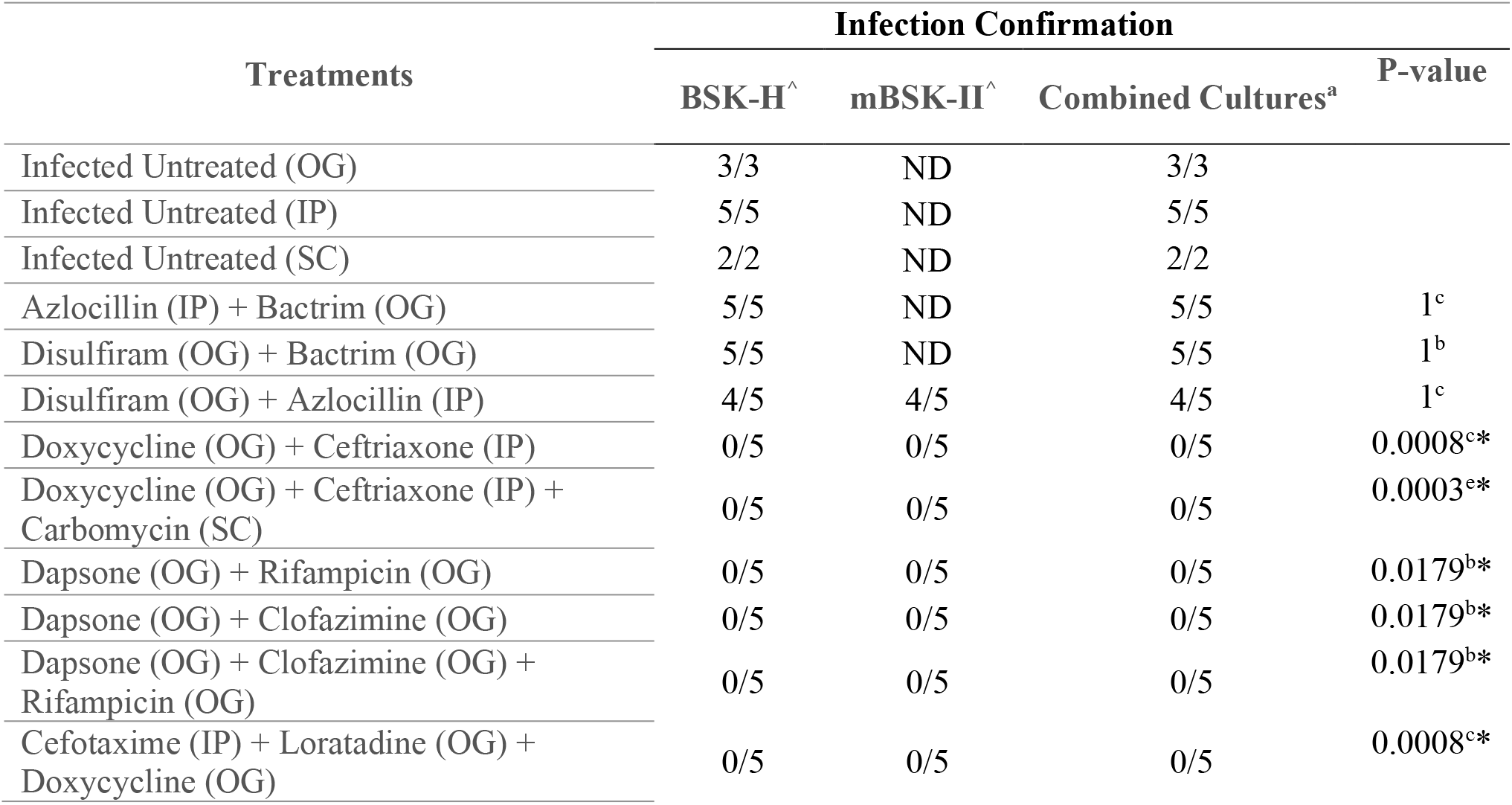
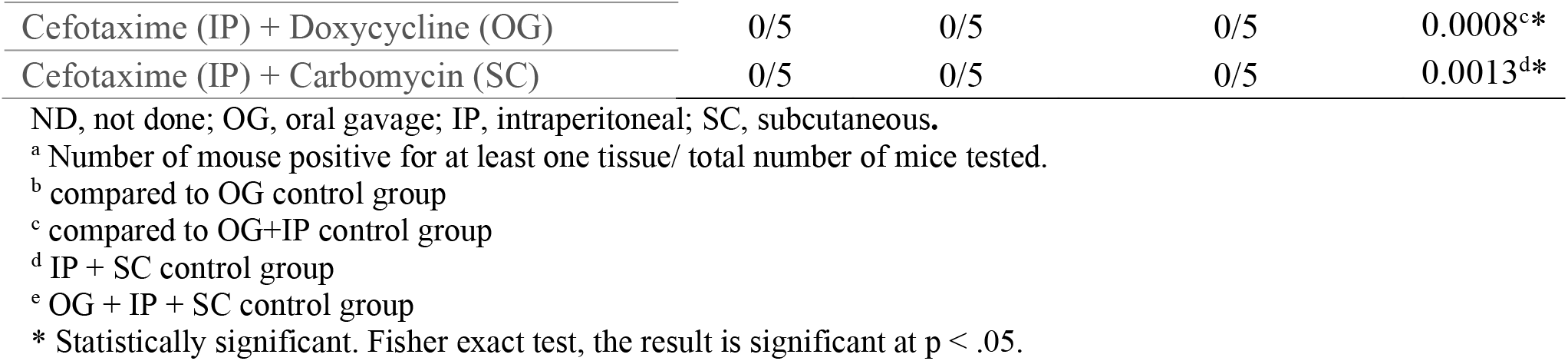
Culture results of tissues in combination therapy.

### RNA not detected in tissues following combination therapy

Groups that were positive for *Bb* infection by the tissue culture assay were excluded from PCR and RT-PCR analyses. *Bb* DNA was detected in at least 1 mouse in each combination group using the *16S* primer and 4 groups (dapsone + rifampicin; dapsone + clofazimine; dapsone + clofazimine + rifampicin; cefotaxime + loratadine + doxycycline) using the *ospC* primer. RT-PCR using *16S* primers did not detect *Bb* in any combination group, indicating complete eradication of viable *Bb. ospC* primers were not used for RT-PCR. All control groups were positive for both primers in PCR and for *16S* in RT-PCR (**Table 6**).

**Table 6:**
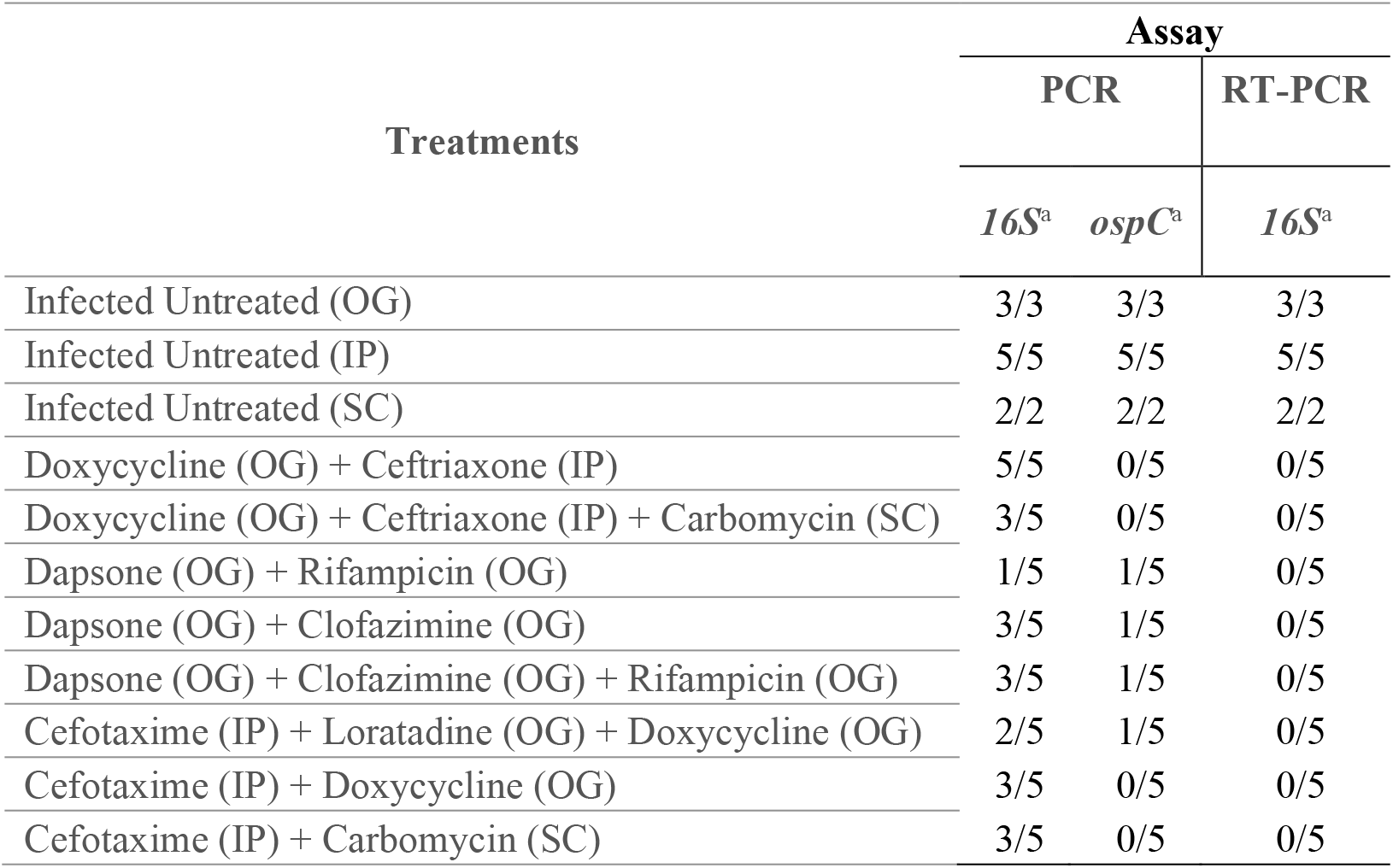
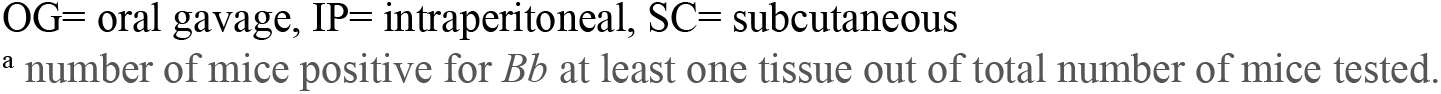
Summary of the result of the molecular detection in tissues.

| Treatments | Assay |  |  |
| --- | --- | --- | --- |
|  | PCR |  | RT-PCR |
|  | *16S*<sup>a</sup> | *ospC*<sup>a</sup> | *16S*<sup>a</sup> |
| Infected Untreated (OG) | 3/3 | 3/3 | 3/3 |
| Infected Untreated (IP) | 5/5 | 5/5 | 5/5 |
| Infected Untreated (SC) | 2/2 | 2/2 | 2/2 |
| Doxycycline (OG) + Ceftriaxone (IP) | 5/5 | 0/5 | 0/5 |
| Doxycycline (OG) + Ceftriaxone (IP) + Carbomycin (SC) | 3/5 | 0/5 | 0/5 |
| Dapsone (OG) + Rifampicin (OG) | 1/5 | 1/5 | 0/5 |
| Dapsone (OG) + Clofazimine (OG) | 3/5 | 1/5 | 0/5 |
| Dapsone (OG) + Clofazimine (OG) + Rifampicin (OG) | 3/5 | 1/5 | 0/5 |
| Cefotaxime (IP) + Loratadine (OG) + Doxycycline (OG) | 2/5 | 1/5 | 0/5 |
| Cefotaxime (IP) + Doxycycline (OG) | 3/5 | 0/5 | 0/5 |
| Cefotaxime (IP) + Carbomycin (SC) | 3/5 | 0/5 | 0/5 |

OG= oral gavage, IP= intraperitoneal, SC= subcutaneous
<sup>a</sup> number of mice positive for *Bb* at least one tissue out of total number of mice tested.

### Culture of xenodiagnostic tick guts from mice treated with combination therapy confirms *Bb* eradication

The same three drug combinations (azlocillin+ Bactrim, disulfiram+ Bactrim, and disulfiram + azlocillin) that were reported as positive for *Bb* as using the tissue culture assay were also positive using the tick mid-gut culture assay. The other treatment groups were negative for *Bb* (*P* ≤.05). The mice in the control groups had a positivity rate of 70%. Results for the positive controls were comparable between BSK-H and mBSK-II media. mBSK-II was not used for OG controls and 3 samples were cultured in mBSK-II compared to 5 in the BSK-H media.

### PCR and RT-PCR confirm eradication of *Bb* infection and viability in xenodiagnostic ticks following combination therapy

Groups that were positive for *Bb* infection in the tissue culture assay were excluded from PCR and RT-PCR analyses. *Bb* DNA was detected using the *16S* primer in all but 2 groups: doxycycline + ceftriaxone + carbomycin and cefotaxime + loratadine + doxycycline. *Bb* DNA was not detected using the *ospA* primer in any of the combination groups.

*Bb* RNA was detected using the *16S* primer in the cefotaxime + carbomycin group. No other RNA was detected in the remaining treated groups using either *16S* or *ospA* primers.

### IFA of xenodiagnostic tick midguts did not detect any spirochetes following combination therapy

Immunofluorescent visualization of tick midguts showed *Bb* spirochetes in 0 out of 5 mice in each combination treatment group (*P* =.0467). Four out of 5 mice in the control group had at least 1 *Bb* spirochete. The IFA was only conducted on ticks in combination groups that were negative in both culture and RT-PCR. Results are summarized in Table 7.

**Table 7:**
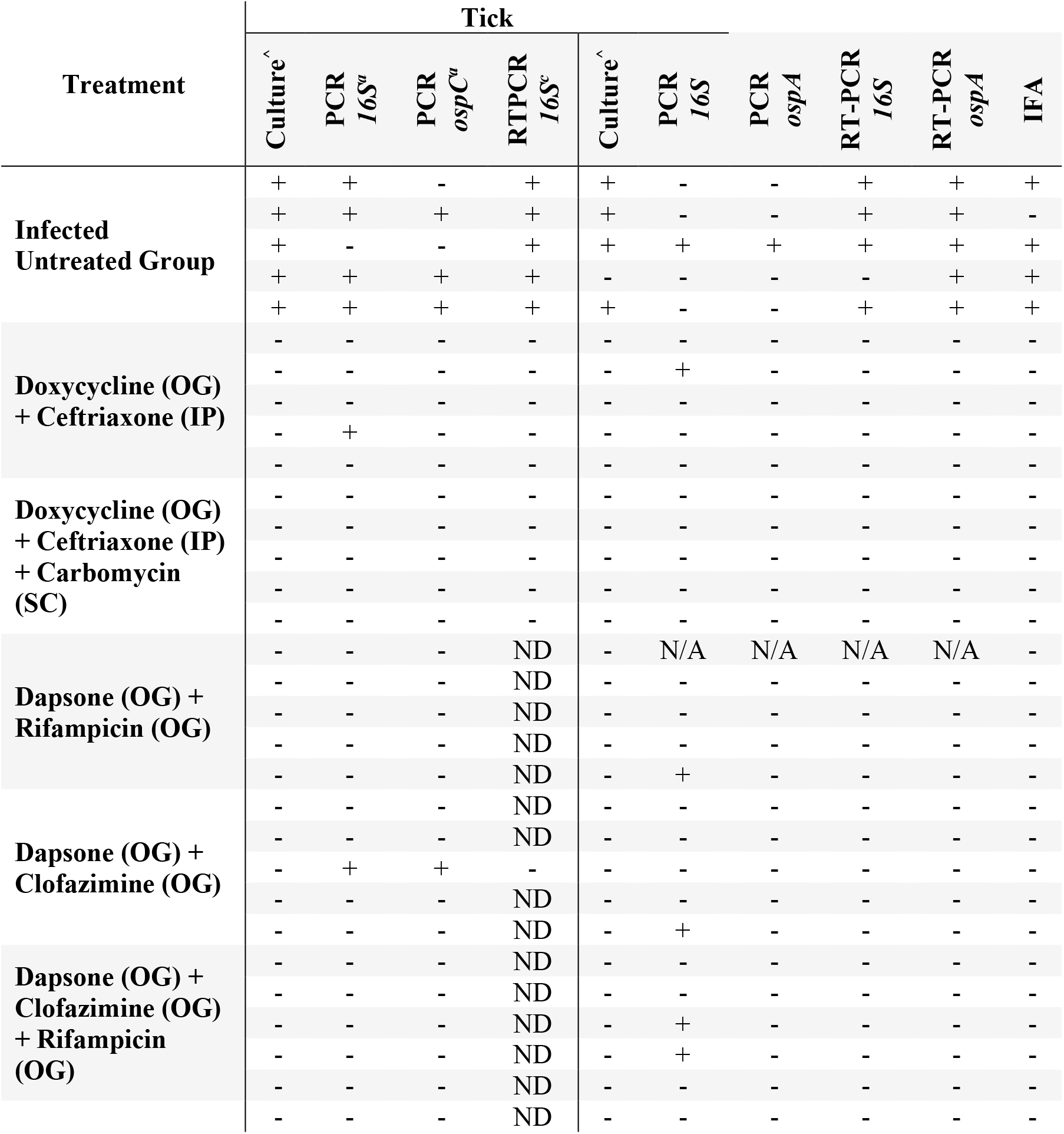
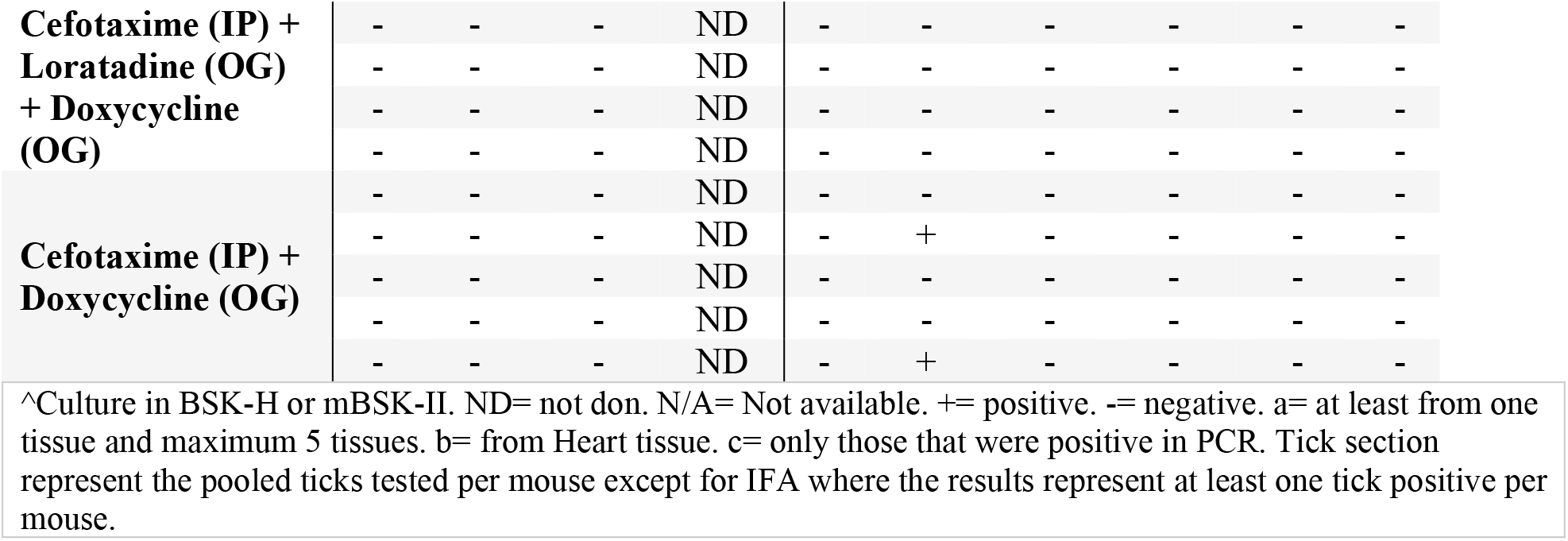
Summary of the test results of the effective treatment groups.

## Discussion

Guidelines dictate that select monotherapies should be used to treat LD, but a subset of patients develop PTLD following treatment. Persistence of small numbers of *Bb* spirochetes following monotherapy in animal models demonstrates that monotherapy does not always eradicate the infection (4). Many other microbial pathogens also cause persistent infection, but some, such as *M. tuberculosis*, are treated with combination therapy to effectively treat the infection (28, 29, 38-43). Unlike *M. tuberculosis* and other persistent pathogens, *Bb* has historically been difficult to culture, making combination therapy studies challenging. The methodology in this study allowed for the cultivation of *Bb* from mouse tissues and xenodiagnostic ticks and detection of viable *Bb* by RT-PCR in mouse tissues and xenodiagnostic ticks. Additionally, PCR and IFA were used to determine *Bb* presence regardless of viability. These techniques provided a platform to test combinations of 2 or 3 FDA-approved monotherapies to treat *Bb* infection.

Two types of media, BSK-H and mBSK-II, were used to culture Bb spirochetes. There was no difference in the number of positive samples between the BSK-H and mBSK-II media, but mBSK-II medium accelerated the growth and the number of spirochetes compared to BSK-H medium.

Our results support past findings that monotherapy does not eradicate *Bb* infection in all mice, leaving a subset of mice with persistent *Bb* infection (15, 16, 18, 44, 45). Viable spirochetes were detected by culturing tissues in 3-5 out of 5 mice in 7 of the monotherapy groups and 1 out of 5 mice for the cefotaxime treatment group. Viability was also confirmed in all monotherapy group xenodiagnostic tick midgut culture and the detection of RNA using RT-PCR in tissues and xenodiagnostic tick midguts. In contrast, several combination therapies were effective; viable spirochetes were not detected in cultured tissues in 8 out of 11 combination therapy groups in all mice. Viable spirochetes were also not detected by xenodiagnostic tick midgut culture or by the presence of RNA in tissues or xenodiagnostic tick midguts for the same combination therapies with the exception of the cefotaxime + carbomycin combination which was positive for *16S* RNA in xenodiagnostic tick midguts. Spirochetes were also not present in tissues when imaged using IFA.

The combination of ceftriaxone + doxycycline may be a good candidate for future investigation. Viable *Bb* was not detected by tissue culture, RT-PCR on tissues, xenodiagnostic tick mid-gut culture, or RT-PCR of xenodiagnostic tick mid-guts. Ceftriaxone and doxycycline are already used as monotherapy for LD treatment and both are well tolerated with favorable safety profiles(19). The addition of carbomycin to the ceftriaxone + doxycycline combination was also examined but the carbomycin did not provide additional benefit.

Cefotaxime + Doxycycline was also effective at eradicating *Bb* infection. Viable *Bb* was not detected by tissue culture, RT-PCR on tissues, xenodiagnostic tick mid-gut culture, or RT-PCR of xenodiagnostic tick mid-guts. The addition of loratadine to the ceftriaxone + doxycycline combination was also examined but the loratadine did not provide additional benefit.

Dapsone in dual or triple combinations with rifampicin, clofazimine, or rifampicin + clofazimine were also effective at eradicating viable *Bb*. The combinations of dapsone with rifampicin, clofazimine or both are already the therapies of choice to treat multidrug-resistant *M. tuberculosis* and leprosy have ample evidence for safety and tolerability in that population (46-49). Furthermore, clinical studies have demonstrated efficacy of dapsone combination therapy in Lyme disease patients with chronic symptoms/PTLD (50-52). The results of this study and well-documented use in humans make this combination attractive for further investigation.

Some combination therapies were not effective at eradicating *Bb*. Three combinations (azlocillin+ Bactrim, disulfiram+ Bactrim, disulfiram + azlocillin) had persistent infection rates similar to those of control mice. Additionally, the mice treated with the cefotaxime + carbomycin combination were negative for *Bb* in tissue and xenodiagnostic tick midgut cultures but *Bb* RNA was detected in the xenodiagnostic tick midguts, indicating viable organisms.

The presence of *Bb* DNA in tissues and xenodiagnostic tick midguts in the absence of *Bb* RNA occurred in mice in most combination therapy groups. DNA in the absence of RNA is likely to indicate that the DNA is from residual non-viable Bb. The extent to which non-viable *Bb* affects the immune response following combination therapy requires further investigation.

Doxycycline and ceftriaxone were not included in this study. They are well studied and are not effective at eradicating the persistent form of *Bb* (17, 19, 45). The purpose of this study was to evaluate the efficacy of monotherapies and combination therapies to eliminate in vivo long-term infection with *Bb*. These results provide evidence that several different combination therapies can completely clear infection in all mice while treatment with monotherapy does not clear the *Bb* infection in some mice. Combination therapies, especially the ones highlighted in this study, need further investigation before conducting human trials. Mice in this study were infected by injection. Future work should examine the efficacy of combination therapy when infection is introduced by tick bite. Infection by tick bite has been previously used in studies using rhesus macaques (17, 19, 20). These combinations should also be tested against other strains of Bb such as N40, JD1, and 297 due to the plasmid variation and genetic diversity among the strains of *Bb* (53), and the loss of several plasmids in the persistent form of spirochetes after antimicrobial treatment in mice (54). Antibiotic susceptibility also varies with strains and isolates (55-57). Additional studies in non-human primates would also provide information important for moving into human clinical trials.

## Materials and Methods

### Murine experiments

The mice used in this study were housed in a pathogen-free animal facility according to animal safety protocol guidelines at the Tulane National Primate Research Center (TNPRC) accredited by the Association for the Assessment and Accreditation of Laboratory Animal Care (AAALAC) International. The Tulane University Institutional Animal Care and Use Committee (IACUC) approved all animal-related protocols. C3H/HeN mice from Charles River labs, aged 6-7 weeks were used in this study. Two to 5 mice were housed per cage and maintained at 25°C with a 12-hour light-dark cycle and had *ad libitum* access to chow and water. Mice were euthanized by CO2 narcosis followed by cervical dislocation and exsanguination by cardiocentesis. Euthanasia was performed in accordance with the guidelines of the American Veterinary Medical Association (AVMA).

### Culture Media

Cultures were grown in BSK-H (Sigma) or mBSK-II medium. mBSK-II medium contains carbohydrate additives including 0.4% each of mannose, maltose, glycerol and n-acetylglucosamine. Phosphomycin, rifampicin, and amphotericin B were added to both medias to prevent fungal growth and bacterial contamination at concentrations recommended in the media protocol (17).

### Ear Biopsy

Ear punch biopsies were collected at days 7 and 14 post-infection using sterile 2mm punches after cleansing the skin with an alcohol wipe. The punches were cultured in BSK-H media and placed in a tri-gas incubator for 5 weeks to confirm infection. Cultures were checked for motile spirochetes once per week.

### *B. burgdorferi* Infection Confirmation

Approximately 1 ×10^6^ *B. burgdorferi* strain B31.5A19, grown from stocks between passages 3-5 were administered by subcutaneous injection. Ear punch biopsies were obtained and cultured at 7 and 14 days post-infection and serum was tested at 0, 21, and 60 days using a 5-antigen test to confirm *Bb* infection (58). Mice were required to be seronegative against the 5 *Bb* antigens at day 0 to be included in the study. Mice were also required to have at least one culture-positive ear biopsy at day 7 or day 14 or be seropositive for 2 or more *Bb* antigens at day 21 or day 60 following inoculation.

Mice were humanely euthanized after the xenodiagnoses (2-3 months after treatment completion and collecting engorged ticks). Skin (one whole ear) was cultured into BSK-II liquid medium and incubated in a microaerophilic chamber, as well as splitting one whole ear for both RNA and DNA extraction. All other tissues (heart, bladder, spleen, tibiotarsal joints) were used for culture and nucleic acid. The growth of *B. burgdorferi* was detected by dark field microscopy with a 50x objective. The RNA and DNA were also detected via RT-PCR or standard PCR.**Other antimicrobial drug treatment**

All drugs (azlocillin, Bactrim, carbomycin, cefotaxime, ceftriaxone, clofazimine, dapsone, disulfiram, doxycycline, loratadine, and rifampicin) used in this study are FDA-approved and were administered at non-toxic levels. Bactrim, Disulfiram, Dapsone, Rifampicin, and Loratadine were administered through oral gavage. Azlocillin and Cefotaxime were administered by intraperitoneal injection. Carbomycin was administered by subcutaneous injection. Drugs were prepared according to the drug specifications.

Oral treatments were given for 28 days and intraperitoneal and subcutaneous treatments were given for 14 days beginning 60 days post-inoculation. Dosage was determined based on weight measured 2 days before treatment.

### Sample Collection for Treatment Efficacy

Blood, tissues (ear, heart, spleen, bladder, and tibiotarsal joint), and tick midguts were collected for this study. Approximately 0.1 ml of blood was collected via retro-orbital bleeding at 0, 14, 21, 60-63, 66-69, 70-73, 80-83, 90-93, 104-107 days and at 6 months post-injection. Blood was also collected at necropsy by cardiac puncture. Serum was eluted from the blood by centrifugation at 6000 rpm for 10 minutes and collecting the supernatant (serum). Serum was preserved at −20 °C until further analysis.

Approximately 2 mm^3^ of tissue from the ear, heart, bladder, tibiotarsal joint (TTJ), and spleen were collected aseptically into PBS at necropsy. Each sample was divided and allocated to either culture medium, a 1.5 ml tube for preservation at −20 °C for PCR, or a 1.5 ml tube with 5 volumes of RNAlater (Qiagen, Valencia, CA) and preserved at -80 °C for RT-PCR.

For xenodiagnoses, six nymphal-stage xenodiagnostic hard ticks, *Ixodes scapularis*, were introduced to each mouse. Engorged ticks detached after 3-6 days of feeding and were kept for 7 to 14 days to allow *Bb* to grow if present. The ticks were washed and crushed individually in 50 μl of PBS using a previously published protocol (17). Tick contents were apportioned to a slide (12 µL), fixed by acetone, and preserved at room temperature for the immunofluorescence assay (IFA). The remaining contents of tick midguts were pooled per mouse and split into 3 equal portions for culture, PCR, and RT-PCR as with the tissue samples.

### *Bb* cultures from tissue

Tissue samples were added to 4 mL of BSK-H or mBSK-II medium, or both. Tick midgut samples were cultured in BSK-H only. Culture tubes were incubated for 5 weeks at 34°C, in the presence of 5% CO_2_ and an influx of N_2_ to produce a microaerophilic environment. The cultures were evaluated for the presence of motile spirochetes every week for 5 weeks using dark-field microscopy.

### Serology Test

Five antigens (OspA, OspC, DbpA, OppA2, and C6) were used concurrently to detect *Bb* antibodies in mouse serum at 0, 21, and 60 days. The Bio-Rad Bio-Plex amine coupling kit (catalog no. 171-406001) protocol for coupling antigens to cytometric beads was slightly modified to include the *C6* peptide and recombinant proteins in the same assay. The modified protocol is described in detail in a previous study (58).

### Determination of Drug Concentration in Serum

Drug concentrations in serum from mice treated with drug(s) or vehicle control were measured using the Modified Kirby-Bauer (MKB) assay on days 7, 14, and /or day 21. In brief, 25 μL of control serum and 25 µL of serum pooled from all mice within a treatment group were pipetted onto a 2-mm diameter plain paper disc (BD) and allowed to dry for 30 minutes. Control standards for each drug were made by dissolving the drugs into serum from untreated mice. Concurrently, a 0.5 McFarland suspension of *Staphylococcus aureus* (ATCC #29213) was made in Tryptic Soy Broth (BD, Franklin Lakes, NJ) and streaked onto Mueller-Hinton agar (MHA) plates (BD). The discs of the standards and treated sera were placed onto the streaked MHA plates and incubated at 37 °C for 18–24 hours. After incubation, the diameter (mm) of the zones of inhibition were measured and a standard curve was generated using the standards measurements. Drug concentration was calculated by comparing the zone of inhibition of the treated sample to the standards (59, 60).

### PCR

DNA was extracted following the protocol of the DNeasy® Blood and Tissue kit (Qiagen, USA). Tissue and tick midgut samples were eluted in 200 μl and 50 μl of elution buffer, respectively. DNA concentration was quantified using a Nanodrop 2000 spectrophotometer (Thermo Fisher Scientific, Wilmington, DE, USA). For tissues, 50-250 ng of template DNA was used, depending on the tissue, and held equal per tissue based on the maximum amount possible from eluted quantities. The PCR was run with 35 cycles of denaturation (94°C, for 30 sec), annealing (57.8°C for *ospC*, 60°C for *ospA*, 62°C for *16S*, for 45 sec) and extension (72°C, for 1 min) using 25 μl of sample. Primers against *16s* and *ospC* were used for tissue samples and primers against *16s* and *ospA* were used for the tick contents. The full sequences of the primers are in **Supplemental Table 1**.

### RT-PCR

Total RNA was extracted from all samples using a RNeasy mini kit (Qiagen, USA) and contaminating DNA was removed using the RNase-free DNase set (Qiagen, USA) according to the manufacturer’s protocol. RNA was reverse-transcribed using a high-capacity cDNA reverse transcription kit (Invitrogen, USA). The same primers used for PCR were used for RT-PCR (except for *ospC* which was not performed for tissues). RT-PCR was performed with the Qiagen® OneStep RT-PCR kit (Qiagen, USA). PCR in the second step was performed as described above.

### Immunofluorescence Assay

Slides of acetone-fixed tick midgut contents were blocked by adding 30uL normal goat serum to each slide (NGS = 10% in PBS + 0.2% Fish Skin Gelatin) and incubated with blocking at 37°C for 1 hour. Slides were washed 3 times with 50 mL of PBS. Slides were then incubated with an anti-OspA mouse monoclonal antibody (CB10; obtained from J. Benach(30)) at a 1:30 dilution in a volume of 30 µL for 1-1.5 hours at 37°C. Slides were again washed 3 times with 50 mL of PBS and incubated with anti-mouse IgG-Alexa 488 secondary antibody (1:1000; ThermoFisher A28175) at 37°C for 1 hour. Slides were washed 3 times with 50 mL of PBS and dried using Kim Wipes while avoiding the treated areas. Before adding coverslips 10-15uL of Anti-Quench was added to each slide. Coverslips were fixed on either side with clear nail polish and allowed to dry. Slides were stored at −20°C until viewing and at 4°C after viewing. The slides were imaged using a Nikon Ti2-E fluorescence microscope to detect the presence of spirochetes.

## Data Analysis

The cut off value for positivity for the 5-antigen test was set for each plate based on the mean MFI of days 0 in that plate for each antigen + the standard deviation of the mean multiplied by 3.

Differences between treated and untreated animals in all culture, PRC, RT-PCR, and IFA assays for tissues and tick midguts were evaluated with Fisher’s exact test (two-tailed). Differences were considered significant at *P* ≤.05.

## Acknowledgements

Funding for this research was provided by the bay Area Lyme Foundation and TNPRC base grant 5 P51OD 011104-56. The authors would like to thank Allie Amick for her contributions to compiling and editing the contents of the manuscript.

